# ER-localized phosphatidylethanolamine synthase plays a conserved role in lipid droplet formation

**DOI:** 10.1101/2021.08.31.458391

**Authors:** Mehmet Oguz Gok, Natalie Ortiz Speer, W. Mike Henne, Jonathan R. Friedman

## Abstract

The asymmetric distribution of phospholipids in membranes is a fundamental principle of cellular compartmentalization and organization. Phosphatidylethanolamine (PE), a nonbilayer phospholipid that contributes to organelle shape and function, is synthesized at several subcellular localizations via semi-redundant pathways. Previously, we demonstrated in the yeast *Saccharomyces cerevisiae* that the PE synthase Psd1, which primarily operates on the mitochondrial inner membrane, is additionally targeted to the endoplasmic reticulum (ER). While ER-localized Psd1 is required to support cellular growth in the absence of redundant pathways, its physiological function at the ER is unclear. We now demonstrate that ER-localized Psd1 sub-localizes on the ER to lipid droplet (LD) attachment sites and further show it is specifically required for normal LD formation. We also find that the role of PSD enzymes in LD formation is conserved in other organisms. Thus, we have identified PSD enzymes as novel regulators of LDs and demonstrate that both mitochondria and LDs in yeast are organized and shaped by the spatial positioning of a single PE synthesis enzyme.

## Introduction

Cellular organelles are defined by the composition of their membrane bilayers, which are composed largely of phospholipids with distinct polar head groups. The biochemical properties of these individual phospholipids give rise to membrane shape, influence protein behavior and complex formation, act as signaling molecules, and contribute to giving organelles their distinct identity and function. Thus, the biosynthesis and spatial organization of these membrane components plays a pivotal role in cellular function.

Phosphatidylethanolamine (PE) is a major phospholipid in cells and has a distinctive shape that contributes to negative membrane curvature (Vance, 2018). PE also serves as a substrate in the biosynthesis of phosphatidylcholine (PC), the most abundant phospholipid in cells. PE is synthesized in multiple cellular locales and via multiple pathways. A common source of PE production is the ER-localized Kennedy pathway, which can be fueled by exogenous ethanolamine or by the breakdown of phospho-sphingosine. Additionally, phosphatidyl decarboxylase (PSD) enzymes utilize phosphatidylserine as a substrate in the production of PE (Di Bartolomeo et al., 2017).

The budding yeast *Saccharomyces cerevisiae* (hereafter yeast) expresses two PSD enzymes, Psd1 and Psd2. Psd2, characterized by its conserved C2 domain, localizes to the endolysosomal system (Gulshan et al., 2010; Kitamura et al., 2002; Trotter & Voelker, 1995). Psd1, meanwhile, is primarily localized to the inner mitochondrial membrane (IMM), where is required for mitochondrial morphology and respiratory function (Chan & McQuibban, 2012; Joshi et al., 2012; Kuroda et al., 2011). Previously, we established that Psd1 can also localize to the ER bilayer, and cells without ER-localized Psd1 have cellular growth defects in the absence of redundant pathways (Friedman et al., 2018). Despite this, the ER-localized Psd1 pool is less abundant than mitochondrial Psd1 and is difficult to detect, particularly in nutrient rich or respiratory-demanding growth conditions (Friedman et al., 2018; Sam et al., 2021). This begs the question of whether the ER-localized enzyme plays a specific cellular function, or if the redundant maintenance of cellular growth is an example of selective pressure to maintain survival in an unusual circumstance.

PSD enzymes are highly conserved from bacteria to humans, though the number of distinct proteins and their subcellular localization are variable depending on the organism. For example, Psd2 is found in other fungi such as fission yeast, but is absent in higher organisms (Luo et al., 2009). Humans have a PSD encoded by the *PISD* gene, which is critical for mitochondrial morphology and function (Steenbergen et al., 2005; Tasseva et al., 2013). Recently, however, human *PISD* was demonstrated to express an alternate isoform (isoform 2) that encodes for an enzyme with a distinct N-terminus (Kumar et al., 2021). When GFP-tagged and overexpressed, this isoform exhibited localization to not only mitochondria, but also to an ER-derived organelle, the lipid droplet (LD). While this work demonstrated that depletion of both isoforms of PISD leads to a defect in the incorporation of oleic acid (OA) into triacylglycerol (TAG), it is unclear whether this is indirectly due to defects in cellular metabolism caused by reduction in mitochondrial function or to a potential specific role of PISD at LDs. Thus, while PISD can target to LDs, it remains unknown whether it plays a physiological role there.

The observation of PISD localization to an ER-associated organelle raised the question of whether ER-localized Psd1 in yeast contributes to LD morphology or function. We therefore examined Psd1 localization in yeast under conditions that promote LD formation utilizing improved fluorescent markers. Here, we show that Psd1 can target on the ER membrane to a subset of LDs, particularly under conditions that promote LD biogenesis. Further, we demonstrate that loss of ER-localized Psd1 causes defects in LD formation and morphology. Finally, utilizing the fission yeast *Schizosaccharomyces pombe*, we show that ER and LD targeting of PSDs is conserved under physiological conditions and that PSD enzymes play a conserved role in contributing to normal LD formation.

## Results

### Psd1 sub-localizes on the ER to a subset of lipid droplets in budding yeast

Previously, we performed confocal fluorescence microscopy of yeast cells expressing EGFP-tagged Psd1 and observed dual localization to the ER and mitochondria (Friedman et al., 2018). To improve our visualization of ER-localized Psd1, we generated a yeast strain with Psd1 chromosomally-tagged with the brighter and more photostable mNeonGreen (mNG). Tagging with mNG did not interfere with Psd1 mitochondrial function, as assessed by growth on carbon sources that require respiration (Fig. 1, figure supplement S1A), nor global cellular function, as assessed by ability to support growth of cells in the absence of other PE-producing pathways (Fig. 1, figure supplement S1B). We imaged cells co-expressing Psd1-mNG, the ER marker mCherry-HDEL, and mitochondrial matrix-targeted mito-TagBFP by confocal fluorescence microscopy and detected ER-localized Psd1 when images were contrast-enhanced to overcome the predominant mitochondrial-localized signal (Fig. 1A). Strikingly, with this improved tag, we routinely detected enrichment of ER-localized Psd1 at discrete focal structures (Fig. 1A, arrows). While the relatively low ER signal and the prevalence of mitochondrial Psd1-mNG signal made a quantitative assessment of bona fide focal structures impossible, a majority of cells displayed localized concentration of Psd1 into foci on the ER membrane.

**Figure 1.**
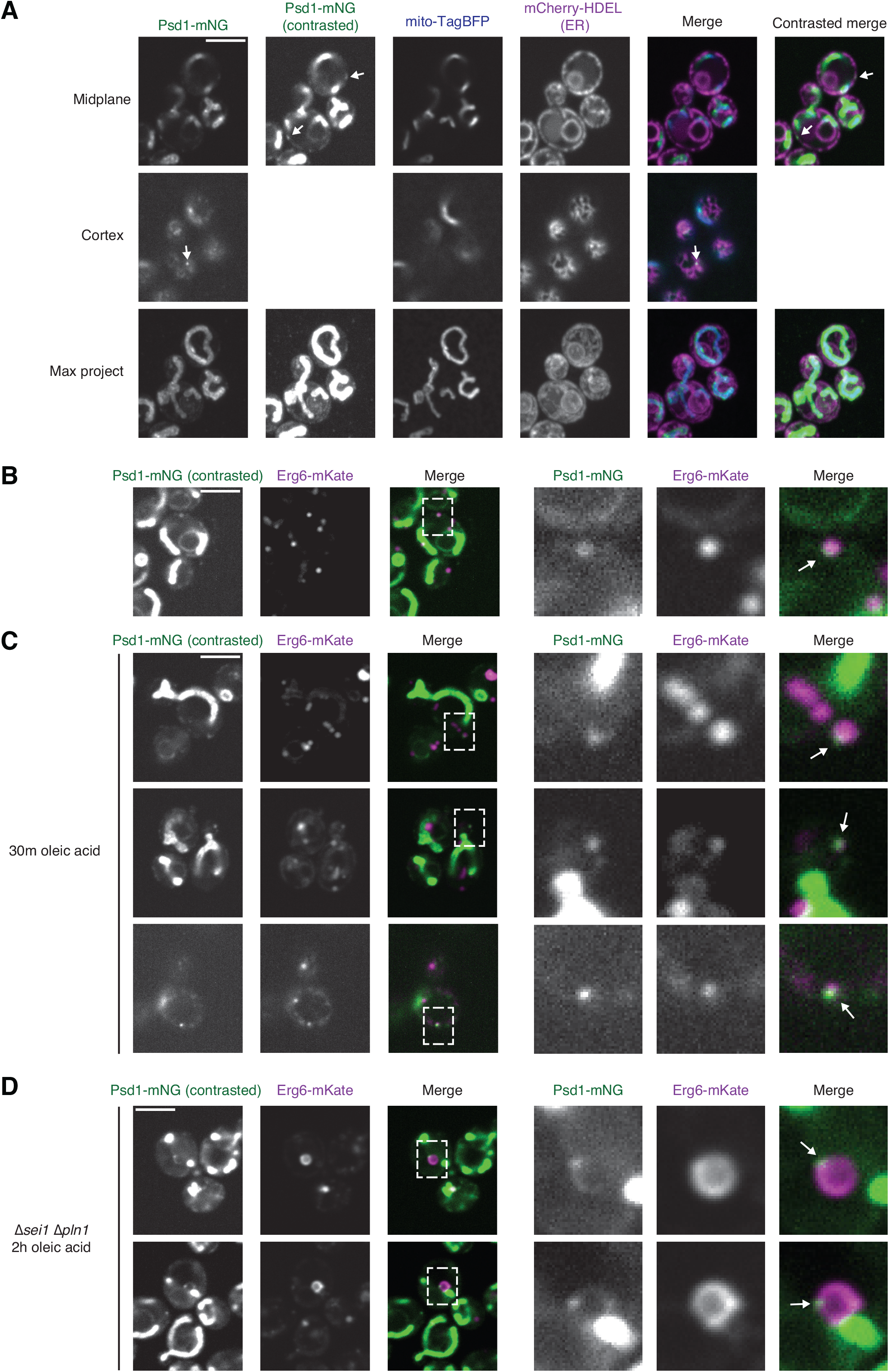
Psd1 sub-localizes on the ER to a subset of lipid droplets in budding yeast. **(A)** Confocal fluorescence microscopy images of wild type cells grown in SD media and co-expressing Psd1-mNeonGreen (mNG; green), mito-TagBFP (blue), and mCherry-HDEL (ER; magenta). Single planes are shown of the cell midplane (top) and cortex (middle); maximum intensity projection is shown at bottom. Arrows mark sites of ER-sublocalized Psd1-mNG puncta. **(B)** Confocal fluorescence microscopy images are shown of a wild type cell co-expressing Psd1-mNG (green) and Erg6-mKate (magenta) and grown in SD media. Images on right are enlarged from boxed region on left. Arrow marks site of Psd1 and Erg6 colocalization. **(C)** As in (B) for cells treated for 30m with 0.2% oleic acid (OA) prior to imaging. **(D)** As in (B) in Δ*sei1*Δ*pln1* cells treated for 2h with 0.2% OA prior to imaging. Psd1-mNG images are shown with non-linear contrast enhancement where indicated to enable visualization of non-mitochondrial signal. Scale bars = 4µm. See also Figure 1 – figure supplement 1.

Recently, work in human cells by Burd and colleagues revealed that the single human PSD, *PISD*, has multiple spliced isoforms that encode for differentially localized proteins (Kumar et al., 2021). When overexpressed and GFP-tagged, a long isoform of PISD exclusively localized to mitochondria, while a shorter isoform localized to both mitochondria and lipid droplets (LDs). As LDs are formed within the ER bilayer (Olzmann & Carvalho, 2019), we reasoned that focal ER-localized Psd1-mNG may target to LDs. To test this, we endogenously tagged the LD marker Erg6 with mKate2 in yeast co-expressing Psd1-mNG and imaged logarithmically growing cells in synthetic defined (SD) media. Under these conditions, we could identify rare examples of unambiguous co-localization between Erg6-mKate and Psd1-mNG (Fig. 2B, arrow). However, we observed many instances with no detectable or inconclusive localization of Psd1 at LDs. Since Psd1 appeared to target only to some LDs and additionally generally localized along the ER bilayer, we reasoned that LD localization of endogenous Psd1 may be transient or occur primarily at a specific stage of LD biogenesis. We thus imaged cells grown in SD media after supplementation for 30 minutes with 0.2% OA to stimulate LD biogenesis. Strikingly, after treatment, Psd1-mNG could routinely be detected at or adjacent to Erg6-mKate labeled LDs (Fig. 1C, arrows).

**Figure 2.**
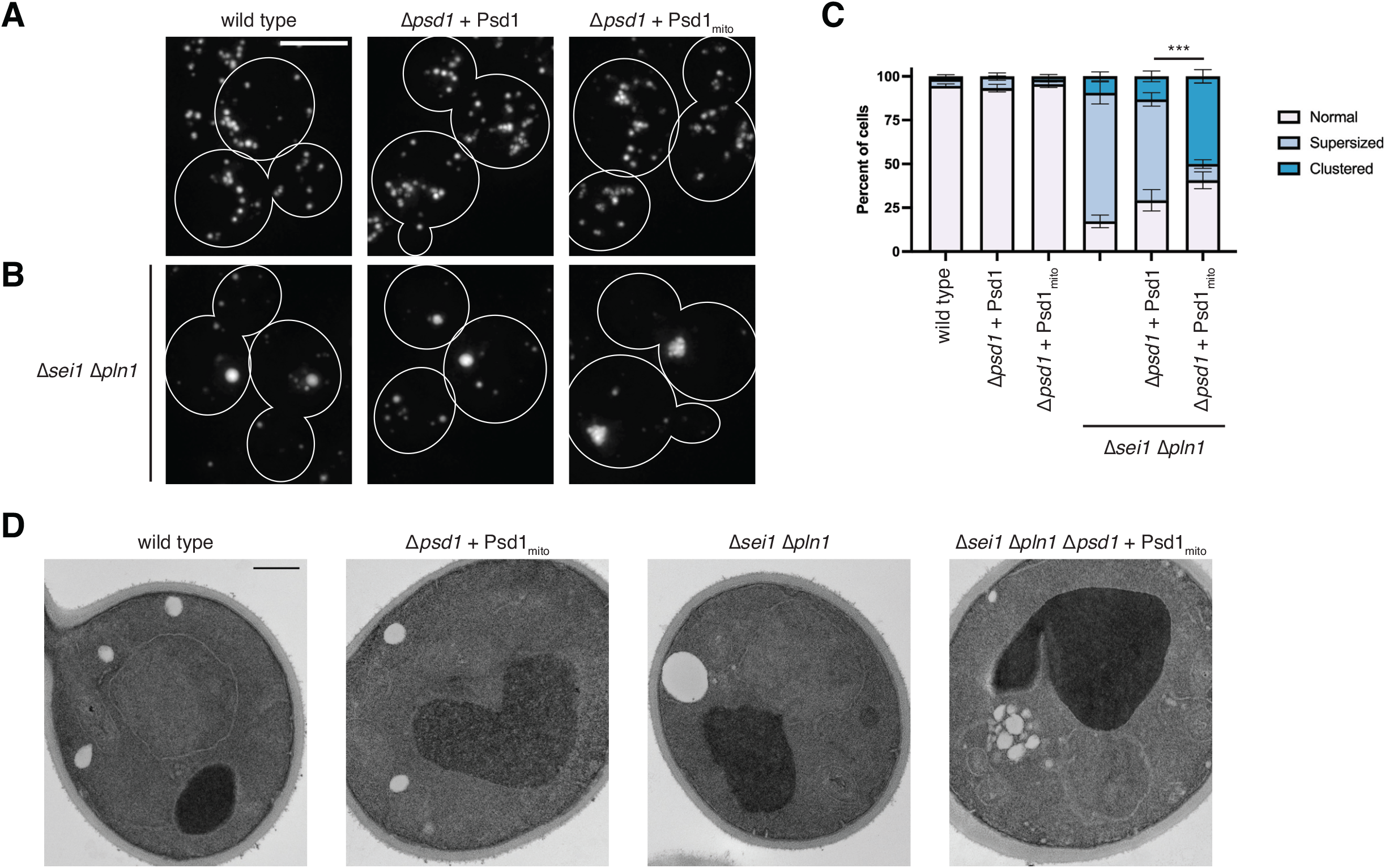
PE generated by ER-localized Psd1 is required for normal lipid droplet morphology. **(A)** Maximum intensity projections of deconvolved epifluorescence microscopy images of LDs in cells from the indicated strain backgrounds grown to exponential phase in SD media, treated for 2h with 0.2% oleate, and stained with the neutral lipid dye MDH. Δ*psd1* cells express wild type Psd1 or Psd1_mito_, where indicated, driven by the native promoter and integrated at the *ura3* locus. Cells are outlined with solid white lines. Scale bars = 4µm. **(B)** As in (A) in Δ*sei1Δpln1* cells. **(C)** A graph of the categorization of LD morphology from cells from (A and B). Data shown are the average of three independent experiments and bars indicate S.E.M. Asterisks (***p<0.001) represent unpaired two-tailed *t* test of supersized LD morphology. See methods for detailed description of categorization. **(D)** Representative electron micrographs from the indicated strains grown to exponential phase in SD and treated for 2h with 0.2% oleate prior to fixation. Scale bars = 500nm.

Because yeast LDs normally appear as diffraction-limited foci by fluorescence microscopy, we sought to alter LD structure in order to more confidently differentiate whether Psd1 targeted to the LD surface, or whether it simply enriched on the ER membrane in close proximity to them. We therefore examined Psd1-mNG localization in the absence of the LD morphogens seipin (Sei1) and Perilipin 1 (Pln1), which leads to the formation of “supersized” LDs (Choudhary et al., 2020; Fei et al., 2008; Gao et al., 2017; Szymanski et al., 2007). We treated Δ*sei1 Δpln1* cells for 2h with 0.2% OA to promote the formation of supersized LDs and, under these conditions, Psd1 localization in proximity to Erg6-labeled LDs was enhanced and we could unambiguously detect Psd1 at the surface of individual LDs. In supersized LDs, Psd1 frequently appeared in focal structures at the LD surface (Fig. 1D, arrows). To a lesser degree, Psd1 also appeared to surround larger LDs (Fig. 1D). Altogether, this indicates that a pool of Psd1 targets to a subset of LDs, particularly under conditions that promote LD biogenesis.

### PE generated by ER-localized Psd1 is required for normal lipid droplet morphology

We next determined whether ER-localized Psd1 contributes to the maintenance of normal LD morphology. We utilized a chimeric allele of Psd1, which replaces the mitochondrial targeting sequence (MTS) and transmembrane domain (TMD) of Psd1 with that of Mic60, another IMM protein (Friedman et al., 2018). This allele, Psd1_mito_, was demonstrated previously to have no detectable ER localization while fully maintaining the mitochondrial functions of Psd1. We labeled LDs with the neutral lipid fluorescent dye monodansylpentane (MDH) in logarithmically growing yeast cells grown in SD media and treated with 0.2% OA for 2h to promote LD biogenesis. As expected, in wild type yeast, LDs appeared as several small punctate structures per cell (Fig. 2A). Similarly, in cells with chromosomally-integrated Psd1 (Δ*psd1* + Psd1) or Psd1_mito_ (Δ*psd1* + Psd1_mito_) driven by the *PSD1* promoter, LDs exhibited normal character, indicating that cells without ER-localized Psd1 did not have a gross LD morphology defect at steady-state (Fig. 2A, 2C).

We reasoned that any defect in LD morphology may be difficult to detect, and therefore examined LD morphology in the *Δsei1* Δ*pln1* background. In these cells, the majority of cells contained LDs with a supersized spherical appearance, which we defined as >0.5µm in diameter (Gao et al., 2017) (Fig. 2B-2C). A small fraction of *Δsei1* Δ*pln1* cells failed to form supersized LDs, and instead exhibited LDs that appeared in a grape-like cluster pattern (9% of cells; Fig. 2C). Unlike cells with wild type Psd1, cells without ER-localized Psd1 (Δ*psd1 +* Psd1_mito_) rarely formed supersized LDs in the Δ*sei1 Δpln1* background; instead they manifested predominantly clustered LDs (50% of cells; Fig. 2B-2C). To investigate the character of LD morphology in greater detail, we performed thin section electron microscopy of cells grown in SD and treated for 2h with 0.2% OA. We regularly observed enlarged LDs in Δ*sei1* Δ*pln1* cells as compared to wild type cells (Fig. 2D). In agreement with fluorescence imaging, Δ*sei1 Δpln1* cells expressing Psd1_mito_ exhibited grape-like LD clusters (Fig. 2D). Together, these data indicate that loss of ER-localized Psd1 exacerbates the LD morphology defect of Δ*sei1 Δpln1* cells and impedes the formation of supersized LDs.

### PE produced by Psd1 is specifically required for the promotion of normal lipid droplet morphology

We next wanted to determine if the contribution of ER-localized Psd1 to LD morphology was specific or mimicked by altering PE production by other means. In addition to Psd1, yeast cells grown in SD media are able to produce PE by two additional pathways (Birner et al., 2001; Gulshan et al., 2010; Trotter & Voelker, 1995). Psd2, while incapable of rescuing the mitochondrial defects caused by loss of Psd1, is able to support cellular growth in its absence. Additionally, Dpl1 is able to generate ethanolamine by degrading phosphorylated sphingoid bases, which can then fuel the ER-localized Kennedy pathway to produce PE. We examined LD morphology using MDH after 2h OA treatment in Δ*sei1 Δpln1* cells that harbored additional deletions of *DPL1*, *PSD2*, or both, and compared each to the ER-deficient Psd1_mito_ strain (Fig. 3A). As before, *Δsei1 Δpln1* cells formed predominantly supersized droplets compared to wild type cells but formed clusters in this background when expressing Psd1_mito_ (Fig. 3A). In contrast, loss of Dpl1, Psd2, or both enzymes in a Δ*sei1 Δpln1* background had a less severe effect on the ability of supersized droplets to form (Fig. 3A).

**Figure 3.**
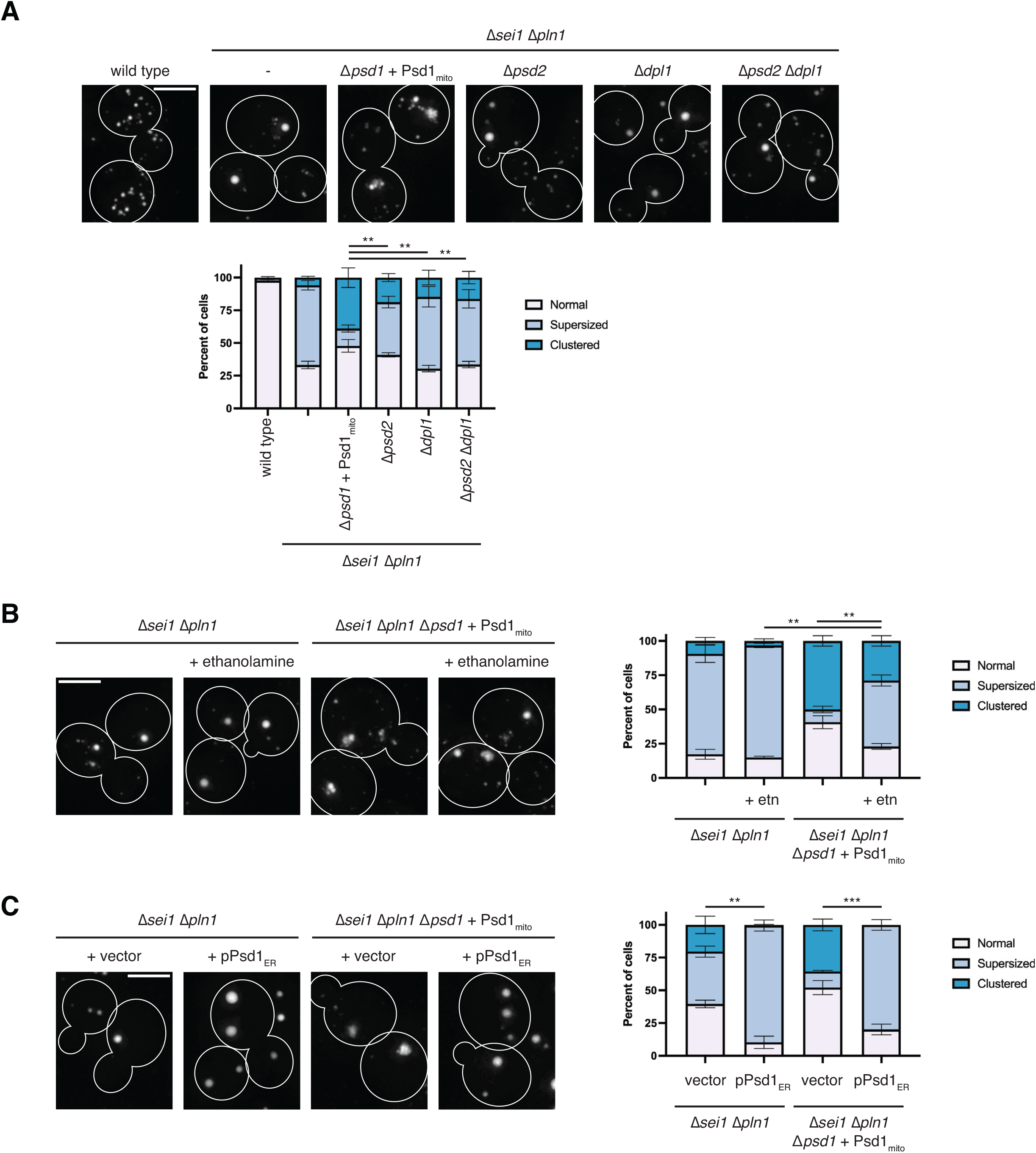
PE produced by Psd1 is specifically required for the promotion of normal lipid droplet morphology. **(A)** Maximum intensity projections of deconvolved epifluorescence microscopy images of LDs in cells from the indicated strain backgrounds grown to exponential phase in SD media, treated for 2h with 0.2% oleate, and stained with MDH. Graph is the categorization of LD morphology from the indicated strains and is the average of three independent experiments. Asterisks (***p<0.001; **p<0.01) represent unpaired two-tailed *t* test of supersized LD morphology. **(B)** As in (A) for the indicated strains. Untreated cells served as controls for multiple experiments and data are redisplayed from Fig. 2C. **(C)** As in (A) for the indicated strains expressing ectopic Psd1 targeted to the ER (pPsd1_ER_). Cells are outlined with solid white lines. Scale bars = 4µm.

Next, we asked whether fueling the Kennedy pathway by supplementing the yeast with exogenous ethanolamine was sufficient to restore the LD morphology of Δ*sei1Δpln1* cells when treated for 2h with OA. Consistent with previous observations (Fei et al., 2011), addition of exogenous ethanolamine alone mildly increased the amount of cells with supersized LDs in a Δ*sei Δpln1* strain (73% of untreated cells vs. 82% of cells treated with ethanolamine; Fig. 3B). Addition of ethanolamine also promoted the ability of Δ*sei1Δpln1* cells with mitochondrial-locked Psd1 to form supersized rather than clustered LDs, however not to the same extent as in yeast expressing wild type Psd1 (48% of ethanolamine treated cells without ER-Psd1 formed supersized LDs; Fig. 3B).

Collectively, these observations suggested that lack of an ER pool of Psd1 contributed to LD morphological defects. To test this, we next assessed whether ectopic expression of ER-localized Psd1 was sufficient to fully rescue the LD morphology of a *Δsei1Δpln1* strain expressing Psd1_mito_. We introduced a plasmid expressing pPsd1_ER_, a Psd1 chimera previously demonstrated to target to the ER using the N-terminal TMD of the ER-resident Sec66 (Friedman et al., 2018). Remarkably, ectopic expression of pPsd1_ER_ qualitatively increased the size and quantitatively increased the prevalence of supersized LDs in Δ*sei1 Δpln1* cells (Fig. 3C). Likewise, expression of pPsd1_ER_ fully rescued the clustered LD morphology defect of cells expressing the Psd1_mito_ chimera (Fig. 3C). Together, these data indicate that Psd1 plays a specific role in contributing to LD morphology at the ER.

### ER-localized Psd1 is required for normal lipid droplet formation

Our observations that ER-localized Psd1 concentrated at a subset of LDs, and that the absence of an ER pool of Psd1 contributed to noticeable differences in LD morphology under conditions that promoted supersized LDs, raised the question of whether Psd1 contributes to LD biogenesis or maturation. In line with this, the presence of non-bilayer lipids like PE on the LD monolayer surface is thought to promote LD fusion and the formation of larger LDs (Chorlay et al., 2019; Fei et al., 2011). To dissect how loss of ER-localized Psd1 impacted LD formation, we modified a system initially developed to induce TAG synthesis and LD formation. This system utilizes a yeast strain in which genes encoding all LD promoting enzymes (Are1, Are2, and Lro1) were deleted except the TAG synthase Dga1, which was placed under control of a carbon-source dependent promoter (Cartwright et al., 2015). We deleted the *PSD1* gene from this strain and reintroduced chromosomally-integrated Psd1 or Psd1_mito_, which were verified to express at equivalent levels to wild type cells and rescue the glycerol growth defects of the Δ*psd1* deficient strain (Figure 4 – figure supplement 1). Given our previous findings that the degree of Psd1 localization to the ER is altered in different carbon sources (Friedman et al., 2018), we engineered the strain to drive expression of Dga1 in the presence of estradiol. Finally, we identified conditions (0.5 nM estradiol) that promoted the formation of nascent LDs in cells grown in SD media on a time scale similar to that observed previously (Gao et al., 2017).

We then asked how loss of ER-localized Psd1 impacted nascent LD formation. As expected, yeast grown in SD media exhibited only very rare LDs by MDH staining (Fig. 4A). Upon treatment with estradiol, LD formation was rapidly induced, with about half of cells having at least one LD after 1h regardless of the presence of wild type Psd1 or Psd1 locked in mitochondria (Psd1_mito_). However, after 2h of treatment, Psd1_mito_ yeast had substantially higher numbers of LDs as assessed by quantifying the number of LDs per cell, or by examining the percentage of cells with 8 or more LDs (Fig. 4B). Indeed, while cells with wild type Psd1 only occasionally formed 8 or more LDs (14% at 4h), nearly half of cells (47% at 4h) with mitochondrial-locked Psd1 formed 8 or more LDs (Fig. 4A-4B). Thus, in cells lacking ER-localized Psd1, LD formation was perturbed, giving rise to excessive small LDs.

**Figure 4.**
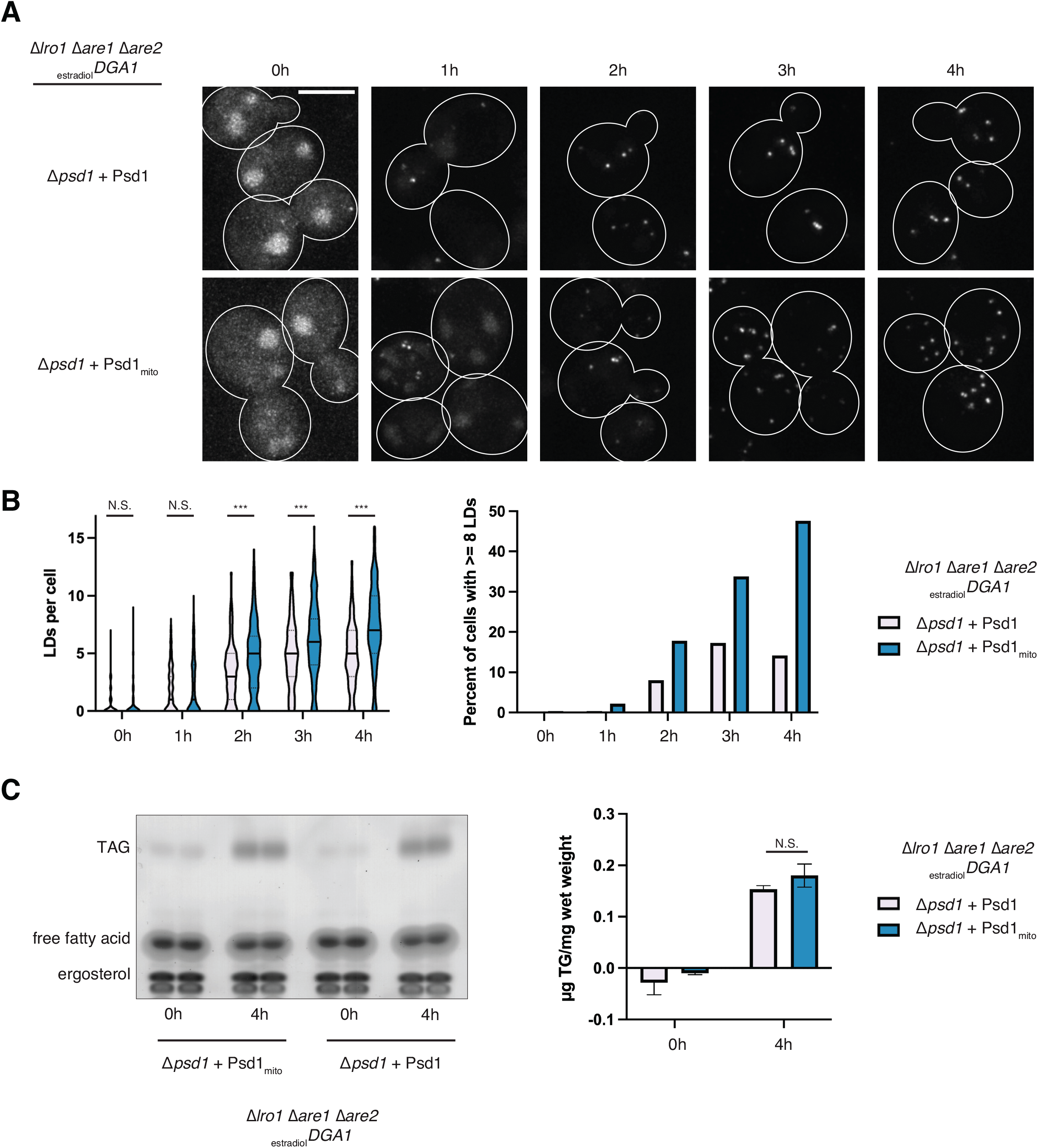
ER-localized Psd1 is required for normal lipid droplet formation. **(A)** Maximum intensity projections of deconvolved epifluorescence microscopy images of MDH-stained LDs in cells expressing wild type Psd1 or Psd1_mito_ where Dga1 expression is controlled by treatment with 0.5nM estradiol in SD media for the indicated times. Cells are outlined with solid white lines. Scale bars = 4µm. **(B)** Graphs of the number of LDs per cell (left) and percentage of cells with 8 or more LDs (right) at the indicated times post-treatment with estradiol from the indicated cells as in (A). Data shown represent a total of 225 cells per strain per timepoint quantified from three independent experiments. Solid lines (left) indicate median and dotted lines indicate upper and lower quartiles. **(C)** Thin layer chromatography analysis of the indicated cells grown as in (A) and treated for the indicated times with 0.5 nM estradiol. Graph (right) displays the amount of TAG/cell weight averaged from three independent experiments. Asterisks (***p<0.001) represent unpaired two-tailed *t* test. N.S. indicates not statistically significant. Bars indicate S.E.M. See also Fig. 4 – figure supplement 1.

To understand the cause of the LD formation defect in cells depleted of ER-localized Psd1, we considered the possibility that the Dga1 enzyme activity may be altered in each strain background. We therefore measured TAG levels in cells with wild type Psd1 and cells with mitochondrial-locked Psd1 by thin layer chromatography. TAG levels were negligible in both strains in untreated cells grown in SD media (Fig. 4C). Importantly, we observed no significant difference in TAG between the strains under conditions where Dga1 expression was induced with estradiol for 4h (Fig. 4C). These data suggest that TAG formation is unaltered in the absence of ER-localized Psd1, and that the increased number of LDs formed upon induced expression of Dga1 are likely due to differences in the coalescence of TAG into nascent LDs.

### Lipid droplet localization of PSD is conserved between budding and fission yeasts

So far, we have determined that in budding yeast, Psd1 is localized to both the mitochondria and to the ER, where it can concentrate near a subset of LDs, particularly under conditions that promote LD biogenesis (Fig. 1). We also have demonstrated that ER-localized Psd1 contributes to normal LD formation (Figs. 2-4). When considering whether this was true of Psd1 orthologs in other organisms, we noted that our previous work identified a glycosylation site on the N-terminus of native Psd1, indicating the ER-localized form of the enzyme must contain a TMD and be integral to the ER membrane (Friedman et al., 2018). In contrast, human PISD is alternatively spliced to produce two isoforms; the canonical long isoform (isoform 1) localizes to the IMM and a shorter isoform (isoform 2) exhibits dual localization (when tagged and overexpressed) at mitochondria and LDs (Kumar et al., 2021). Importantly, the LD-localized isoform in human cells does not contain a predicted TMD and likely targets to LDs via an amphipathic *α*-helix (Kumar et al., 2021). Additionally, while PISD-depleted cells exhibit a defect in incorporation of radiolabeled oleate into TAG, it is unclear whether the LD-targeted form of PISD plays a direct role in LD biogenesis as, to date, no experiments have selectively depleted the LD form of the enzyme (Kumar et al., 2021). Thus, while yeast Psd1 and human PISD likely target to LDs via distinct mechanisms, whether PSD plays a conserved role in contributing to LD formation is an open question.

To investigate whether additional Psd1 orthologs localize to LDs, we evaluated the presence of PSD in other model systems. Remarkably, the fission yeast *Schizosaccharomyces pombe (Sp)* has three distinct PSD enzymes. As in budding yeast, *S. pombe* cells require a source of PE production for cell growth, and loss of all three PSD enzymes is lethal unless cells are supplied with exogenous ethanolamine; however, each enzyme alone is also sufficient to support life (Luo et al., 2009). Psd1 is conserved in *S. pombe* and also named Psd1 *(hereafter Sp*Psd1 for clarity) and contains a computationally predicted MTS and TMD, akin to isoform 1 of human PISD (Fig. 5A). Additionally, Psd2, which can be identified by its C2 domain, is not conserved in humans, but is found in *S. pombe* and named Sp*Psd3*. Finally, *S. pombe* has an additional PSD enzyme, *Sp*Psd2, which despite its name, is distinct from *S. cerevisiae* Psd2. *Sp*Psd2 has no predicted TMD and the presence of an MTS is ambiguous (Fukasawa et al., 2015). Interestingly, in genome-wide analysis of overexpressed GFP-tagged *S. pombe* open reading frames, *Sp*Psd2 was annotated to target to both mitochondria and the nuclear envelope (Matsuyama et al., 2006). This raised the question of whether *Sp*Psd2 may exhibit similar behaviors to the ER-targeted Psd1 of *S. cerevisiae* and isoform 2 of human PISD.

**Figure 5.**
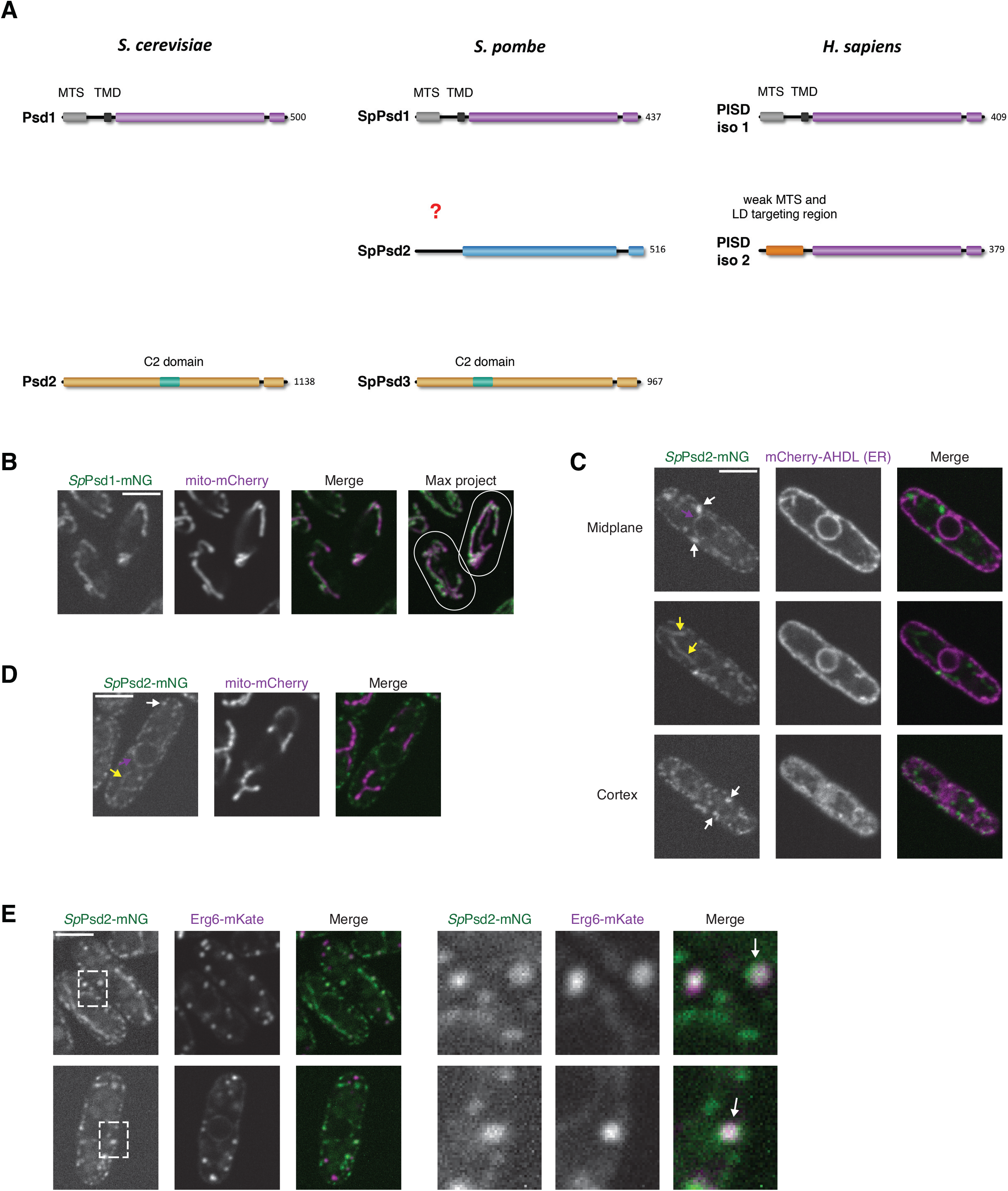
Lipid droplet localization of PSD is conserved between budding and fission yeasts. **(A)** Schematic depicting conservation of PSD enzymes between *S. cerevisiae*, *S. pombe*, and *H. sapiens*. **(B)** Confocal fluorescence microscopy images of wild type fission yeast cells co-expressing *Sp*Psd1-mNG (green) and mito-mCherry (magenta) and grown in EMM media. Single planes are shown except where indicated. Cells are outlined with white lines. **(C-E)** As in (B) for cells expressing *Sp*Psd2-mNG (green) and (C) mCherry-AHDL (ER; magenta), (D) mito-mCherry (magenta), or (E) Erg6-mKate (magenta). Dashed boxes (E) indicate area of enlargement on right. Magenta arrows mark localization of *Sp*Psd2 to the ER, yellow arrows mark localization of *Sp*Psd2 to mitochondria, and white arrows mark localization of *Sp*Psd2 to LDs. Scale bars = 4µm.

We examined localization of *Sp*Psd1 and *Sp*Psd2 by chromosomally tagging mNeonGreen at the C-terminus of both genes in *S. pombe*. Consistent with predicted targeting to the IMM, *Sp*Psd1-mNG localized to mitochondria, which we verified by co-expressing mito-mCherry and imaging cells grown in the synthetic glucose medium EMM (Fig. 5B). In contrast to Psd1 from *S. cerevisiae*, we did not detect obvious ER targeting of *Sp*Psd1. We next examined the localization of *Sp*Psd2-mNG. Remarkably, *Sp*Psd2 appeared to localize to multiple distinct subcellular compartments. In confocal sections taken through the cell midplane, we observed targeting to the nuclear envelope and in the cell periphery, which we verified by coexpressing the lumenal ER marker mCherry-AHDL (Fig. 5C). In other sections of the cells, we noticed apparent mitochondrial targeting of *Sp*Psd2, which we verified by co-expression of mito-mCherry (Fig. 5C-5D). However, we frequently observed a pool of *Sp*Psd2 that appeared in focal structures adjacent to the ER membrane, even without supplementation with OA (Fig. 5C). To determine if these foci were co-localized with LD markers, we chromosomally tagged the C-terminus of the *S. pombe* Erg6 ortholog with mKate (Meyers et al., 2017). Indeed, Erg6-labeled LDs frequently co-localized with focal *Sp*Psd2, indicating that LD targeting of PSD enzymes is conserved in three distinct model organisms (Fig. 5E).

### Loss of spPsd2 impacts LD morphology in fission yeast

We next dissected the functional roles of the different PSD enzymes in *S. pombe*. As both *Sp*Psd1 and *Sp*Psd2 were observed to localize to mitochondria, we asked whether deletion of each affected the ability of cells to grow on nutrient rich media (YES) supplemented with glucose, which does not require mitochondrial respiration, or media with a non-fermentable carbon source (ethanol/glycerol). Similar to in *S. cerevisiae*, *Sp* Δ*psd1* cell growth was severely affected specifically on respiration-requiring media (Fig. 6A). In contrast, *Sp Δpsd2* cells had no observable growth deficiency on either carbon source. We then asked if combined loss of both Δ*psd1* and Δ*psd2* in *S. pombe* exhibited a more severe growth defect but observed no additive effect of loss of *Sp*Psd2 (Fig. 6A). These data indicate that mitochondrial targeting of *Sp*Psd2 contributes negligibly to the respiratory dependent growth of *S. pombe*, while *Sp*Psd1 primarily contributes to respiratory function.

**Figure 6.**
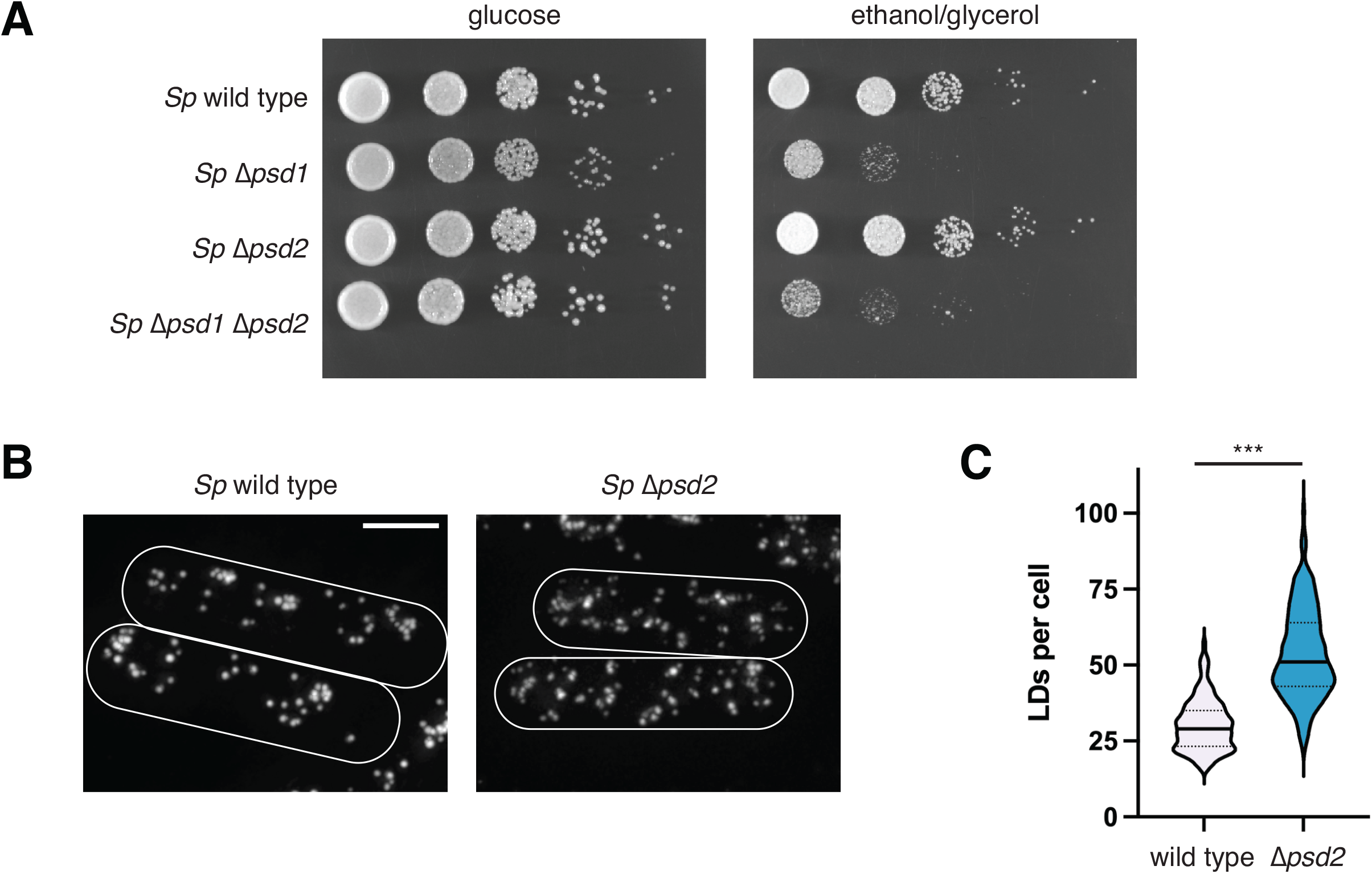
Loss of *Sp*Psd2 impacts LD morphology in fission yeast. **(A)** Serial dilutions of the indicated fission yeast cells plated on YES media containing glucose (left) or the non-fermentable carbon source ethanol/glycerol (right). **(B)** Maximum intensity projections of deconvolved epifluorescence microscopy images of LDs in cells from the indicated strains grown to exponential phase in EMM media, treated for 4h with 0.2% oleate, and stained with MDH. Cells are outlined with solid white lines. Scale bars = 4µm. **(C)** Graph of the number of LDs per cell from the indicated strains as in (B). Data shown represent a total of at least 260 cells per strain quantified from three independent experiments. Solid lines indicate median and dotted lines indicate upper and lower quartiles. Asterisks (***p<0.001) represent results of unpaired two-tailed *t* test.

Given that deletion of *S. pombe* Psd2 did not appreciably affect mitochondrial function, we asked how LD morphology was affected in cells. We stained LDs with MDH in logarithmically growing cells in the synthetic glucose medium EMM after treatment for 4h with 0.2% OA. Remarkably, LDs were far more prevalent in Δ*psd2* cells, with approximately twice as many LDs per cell as compared to wild type cells (Fig. 2B-2C). While this defect was more severe than loss of ER-localized Psd1 in *S. cerevisiae* as it was observed at steady state rather than during induced LD biogenesis, these data are consistent with the increased LD number observed during induced nascent LD biogenesis in budding yeast. Together, these observations suggest a conserved role for ER/LD-localized PSD in promoting the formation of a normal number of LDs per cell.

## Discussion

LDs are surrounded by a phospholipid monolayer that influences protein targeting to the LD surface, but how distinct phospholipids contribute to the biogenesis and maturation of LDs is poorly understood. We have determined that PE-synthesizing PSD enzymes play a conserved role in LD formation in budding and fission yeasts. In budding yeast, the PSD Psd1 localizes both to mitochondria, where it is critical for respiratory function, and to the ER, where it can be observed to concentrate in close proximity to a subset of LDs, likely at the site of connection between LDs and the ER network. Further, we demonstrate that ER-localized Psd1 is required for proper LD formation. Our data indicate a specific functional role of Psd1 in organelle morphogenesis that cannot be completely compensated for by other sources of PE, including exogenous ethanolamine utilized by the Kennedy pathway. We also find that in fission yeast, the PSD *Sp*Psd2 localizes to three distinct organelles: mitochondria, the ER, and LDs. We show that the primary mitochondrial PSD function is performed by *Sp*Psd1, while *Sp*Psd2 is functionally required for the formation of an appropriate number of LDs per cell. Combined with recent observations that humans have a PSD that can target to LDs (Kumar et al., 2021), our work demonstrates that a primary and conserved function of ER/LD-localized PSD is to spatially contribute to organelle biogenesis.

Due to the technical limitations of detecting the minor amount of yeast Psd1 at the ER relative to the abundant mitochondrial pool, it was not possible for us to temporally link Psd1 concentration in proximity to LDs to the biogenesis of the organelle. However, several observations indicate that Psd1 likely transiently localizes to and generates PE at the site of LD formation. First, we rarely detect ER-localized Psd1 foci that co-localize with Erg6 except upon brief stimulation of LD formation with oleate. Second, in the absence of seipin and Pln1, where LD morphology is enlarged, Psd1 clearly enriches at puncta on LDs, likely where it connects with the ER membrane. Finally, while loss of ER-localized Psd1 does not cause a noticeable LD morphology defect at steady state except in Δ*sei1* Δ*pln1* cells, it does cause a significant increase in LD copy number during *de novo* LD biogenesis. Unlike budding yeast Psd1, fission yeast *Sp*Psd2 does not have a predicted TMD, potentially revealing it targets to LDs similarly to human PISD isoform 2, which can be easily observed in a ring around the entire LD and likely localizes via an amphipathic helix (Kumar et al., 2021). However, the increased LD number in *S. pombe Δpsd2* cells implies either enhanced LD formation events as observed in *S. cerevisiae* or a decrease in LD-LD fusion. Thus, a common role of PSDs may be to generate PE to promote TAG coalescence and/or LD growth (Fig. 7). Loss of this local PE pool in the ER network may thus give rise to larger numbers of smaller nascent LDs, perhaps by altering membrane composition leading to premature LD budding or dysregulated emergence (Adeyo et al., 2011; Ben M’barek et al., 2017; Choudhary et al., 2018). Additionally, in yeast genetic backgrounds that support the formation of supersized LDs (such as *Δsei1 Δpln1*), the absence of ER-localized Psd1 may reduce the PE levels on the LD surface monolayer, attenuating the ability of small LDs to fuse with one another and generate supersized LDs (Fei et al., 2011). A non-mutually exclusive possibility is that altered PC/PE ratios at the ER in PSD-deficient cells influences LD protein targeting, which feeds back into influencing LD biogenesis or expansion (Caillon et al., 2020).

**Figure 7.**
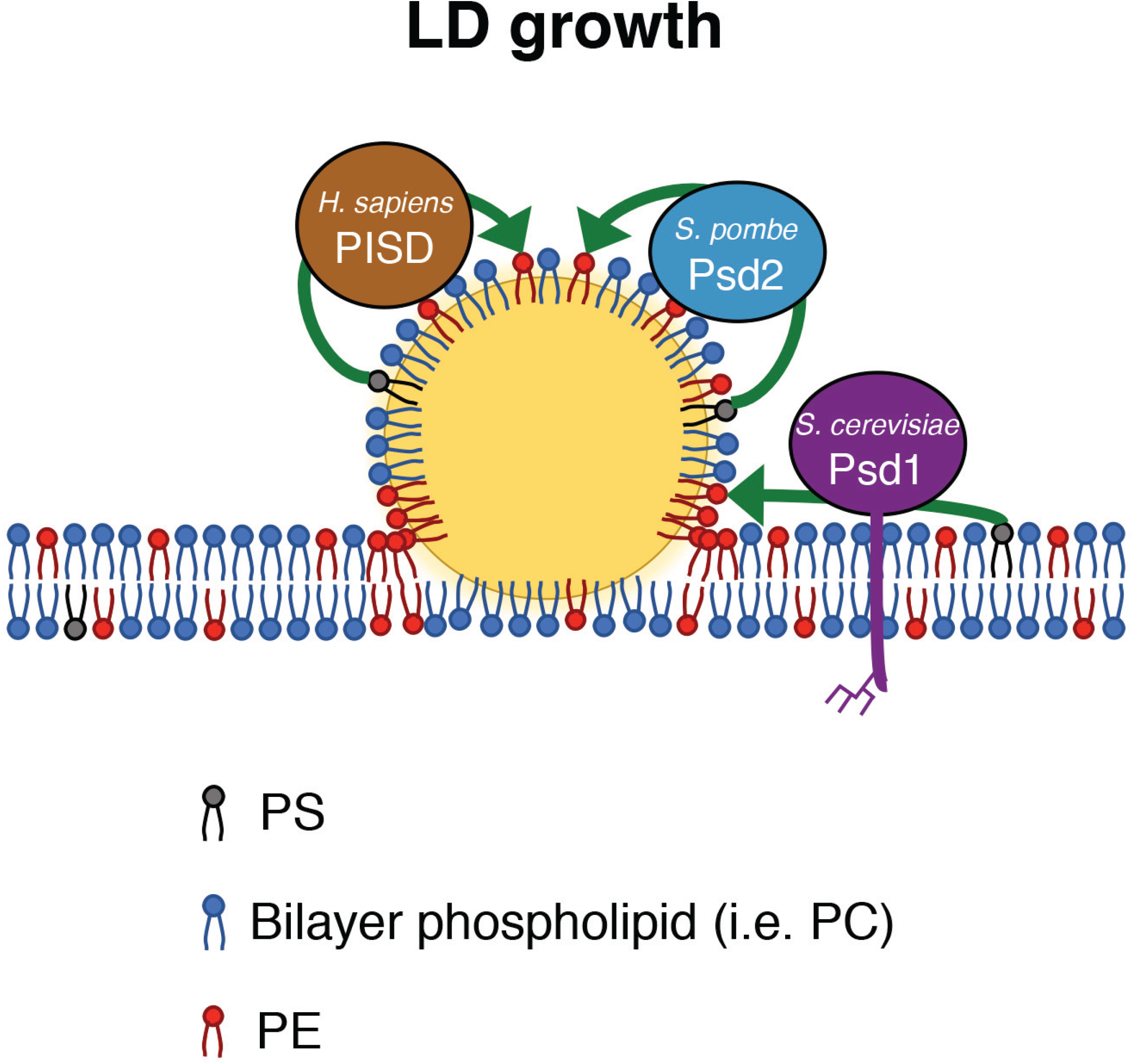
Model for the role of PSD enzymes at sites of LD formation and growth. In yeast, Psd1 concentrates at discrete positions on the ER membrane during LD biogenesis and expansion. Psd1 and its homolog, *S. pombe* Psd2, maintain the appropriate number of LDs during their biogenesis by contributing locally to the production of PE from PS (arrows). Yeast Psd1 is integral to the ER membrane, where it is glycosylated, while *S. pombe* Psd2 and human PISD are soluble and may target directly to the LD surface. Our data are consistent with the model that PE produced locally by PSD enzymes helps facilitate, in conjunction with known LD promoting factors such as seipin, either LD formation and/or later stages of LD maturation, such as fusion. This may occur by the production and concentration of PE locally at the LD neck, a site of negative membrane curvature. PSD enzymes may also contribute to LD growth and LD-LD fusion by producing PE on the LD surface itself. Additionally, by altering the LD surface phospholipid composition, LD behavior may be indirectly affected by impacting protein targeting to the organelle.

A common feature between Psd1, *Sp*Psd2, and the short isoform of human PISD is their localization to multiple organelles. In budding yeast, the mitochondrial Psd1 is integral to the inner membrane and is required for respiratory function. We previously determined that different metabolic growth conditions alter the degree of ER localization of Psd1 (Friedman et al., 2018). Given the roles of LDs and mitochondria in cellular metabolism, an outstanding question is how the cell controls the amount of enzyme targeted to each compartment. Interestingly, in human cells, the relative amount of mitochondrial-versus LD-targeted PISD isoform 2 changes dependent on metabolic growth conditions (Kumar et al., 2021), suggesting a conserved mechanism may exist to regulate differential targeting. In addition to mitochondrial versus ER/LD targeting, it also remains to be determined how yeast Psd1 partitions on the ER and concentrates at LDs. Previously, LD targeting of some proteins was shown to protect against their degradation on the ER (Ruggiano et al., 2016), leaving open the possibility that a similar regulatory mechanism controls Psd1 distribution.

A key difference between budding and fission yeasts is that mitochondrial targeting of *Sp*Psd2 is redundant with *Sp*Psd1. We found that *Sp*Psd1 is required for respiratory growth, while *Sp*Psd2 was dispensable, even in the absence of *Sp*Psd1. These data raise the question of what, if any, physiological role *Sp*Psd2 plays at the mitochondria. Unlike *Sp*Psd1, *Sp*Psd2 has no predicted TMD domain, though it likely contains an MTS and localizes to the mitochondrial matrix. Interestingly, the short isoform of human PISD also localizes to mitochondria via an MTS (Kumar et al., 2021), raising the possibility that these enzymes may play a specialized role in generating PE on the matrix-facing leaflet of the IMM. Alternatively, their import into mitochondria may serve to sequester excess enzyme from activity at the ER/LD membrane.

While we have now demonstrated that ER-localized PSD enzymes play a conserved role in LD formation, PSD can also be observed to localize in general to the ER membrane in three different model systems. An outstanding question is whether PSD has other roles at the ER, for example under circumstances that promote biogenesis of peroxisomes, which can also be formed at LD biogenesis sites (Joshi et al., 2018), or during UPR-mediated ER expansion (Schuck et al., 2009). It also is unclear to what extent that ER-localized Psd1 generates PE that is ultimately converted to PC, or whether mitochondrial-produced PE is the major substrate for PC formation and ER-localized Psd1 is only active under specific growth conditions. While the amount of ER-localized PSD is relatively minor compared to the mitochondrial enzyme, at least in budding yeast, it is now clear that it plays a fundamentally important role in organelle biogenesis.

## Acknowledgements

We thank Madeleine Vaughn for technical contributions. We thank Joel Goodman for helpful discussions and for kindly providing yeast strains, Laura Lackner for kindly providing plasmids, and Steve Claypool for generously providing Psd1 antibody. The UT Southwestern Live Cell Imaging Facility, which is supported in part by P30CA142543, provided access to the Nikon spinning disk microscope (purchased with 1S10OD028630-01 to KLP) and deconvolution software. The UT Southwestern Electron Microscopy facility prepared samples for analysis. This work was supported by grants from the NIH (R00HL133372 and R35GM137894 to J.R.F; R35GM119768 and R01DK126887 to W.M.H), the Welch Foundation (I-1873 to W.M.H.), and the UT Southwestern Endowed Scholars Program to J.R.F. and W.M.H. N.O.S. was supported by NIH T32GM007062.

## Author contributions

Conceptualization, J.R.F. and W.M.H.

Investigation, M.O.G., N.O.S., and J.R.F.

Formal analysis, M.O.G., N.O.S., and J.R.F.

Writing – Original Draft, J.R.F.

Writing – Review and Editing, J.R.F., M.O.G., and W.M.H.

## Materials and Methods

### Strains, plasmids, and media

All *S. cerevisiae* strains were constructed in the W303 genetic background (*ade2-1; leu2-3; his3-11, 15; trp1-1; ura3-1; can1-100*). Wild type haploid *S. pombe* was a kind gift of Jeffrey Pleiss.

All *S. cerevisiae* strains were grown as indicated in YPD (1% yeast extract, 2% peptone, 2% glucose), YPEG (1% yeast extract, 2% peptone, 3% glycerol, 3% ethanol), SD (2% glucose, 0.7% yeast nitrogen base, amino acids) at 30°C. All *S. pombe* strains were grown as indicated in YES (0.5% yeast extract, 3% glucose, 225 mg/L adenine, 225mg/L leucine, 225 mg/L histidine, 225 mg/L uracil, 225 mg/L lysine) or EMM (Sunrise Science; supplemented with 225 mg/L adenine, 225mg/L leucine, 225 mg/L histidine, 225 mg/L uracil, 225 mg/L lysine) at 30°C. To test respiratory growth, YES media was prepared with 3% glycerol, 3% ethanol in place of glucose. Where indicated, cells were treated with 0.2% v/v oleic acid (Sigma O1008), 10mM ethanolamine (pH5.6, ACROS Organics 149582500), or 0.5 nM *β*-estradiol (Sigma/Calbiochem 3301).

All deletions were made using PCR-based homologous recombination replacing the entire ORF with the NatMX6 or HphMX6 cassette from pFA6a-series plasmids using lithium acetate transformation (Longtine et al., 1998). C-terminal protein fusions were integrated at the endogenous loci using pYLB10(mNeonGreen(yeast optimized)-hphMX6) (Arguello-Miranda et al., 2018), pYLB9(mNeonGreen(yeast optimized)-NatMX6 (Wood et al., 2020), pFA6a-mNeonGreen-KanMX6 (see below), pFA6a-link-yomKate2-SpHis5 (Lee et al., 2013), and pFA6a-mKate2-NatMX6 (see below). The Δ*sei1*Δ*pln1* strain was a kind gift of Joel Goodman. Combinations of multiple tags and/or deletions were generated by back-crossing and tetrad dissection and/or by serial PCR-based homologous recombination.

To examine lipid droplet biogenesis in *S. cerevisiae, Δpsd1*::NatMX (Friedman et al., 2018) was introduced into the *3KO(_GAL_DGA1)* strain (Cartwright et al., 2015) by crossing and tetrad dissection. pRS306 Psd1-HygR and pRS306 Psd1_mito_-HygR (see below) were linearized and introduced into that strain and expression was verified by Western analysis (see Figure 4 – figure supplement 1). To control *DGA1* expression with estradiol, pAGL-KanMX6 (see below) was linearized and replaced the *leu2-3* locus.

pFA6-mNeonGreen-KanMX6 was generated by digesting mNeonGreen from pFA6a-mNeonGreen-HygR and cloning into the PacI/AscI sites of pKT127 (pFA6a-yEGFP-Kan) (Sheff & Thorn, 2004). pFA6a-mKate2-NatMX was generated by cloning the NatMX6 cassette from pFA6a-NatMX6 digested with BglII/EcoRI and cloning into the BglII/EcoRI sites of pFA6a-yomKate2-SpHis5.

pRS306 Psd1-HygR and pRS306 Psd1mito-HygR were generated by digesting the HphMX6 cassette from pFA6a-HphMX6 with SacI/NotI and ligating it into the SacI/NotI sites in pRS306 Psd1 and pRS306 Psd1mito, respectively (Friedman et al., 2018). pAGL-KanMX6 was generated by PCR amplifying the KanMX6 cassette from pFA6a-KanMX6 and cloning into pAGL (Veatch et al., 2009) digested with AscI/BsmI by isothermal assembly.

To reintroduce pPsd1_ER_ into cells, pRS314 Psd1_ER_ was generated by cloning the Psd1ER cassette from pRS306 Psd1_ER_ (Friedman et al., 2018) with NotI/KpnI and ligating it into the NotI/KpnI sites of pRS314 (Sikorski & Hieter, 1989).

To visualize the ER in *S. cerevisiae* cells, pRS305 mCherry-HDEL (Friedman et al., 2018) was linearized with EcoRV and integrated into the *leu2-3* locus. To visualize mitochondria in *S. cerevisiae* cells, pVT100-mitoTagBFP (Friedman et al., 2015) was used.

To visualize the ER in *S. pombe* cells, pAV0764 (mCherry-AHDL; kindly provided by Sophie Martin - Addgene # 133520) was linearized with BstZ17I and integrated into the *lys3* locus and selected for with blasticidin (Vjestica et al., 2020). To visualize mitochondria in *S. pombe* cells, mito-mCherry (mitoRED::Hyg; (Kraft & Lackner, 2019)) was linearized with NotI and introduced into the *leu1* locus and selected for with hygromycin.

### Cell growth analysis

For analysis of growth on glucose versus ethanol/glycerol solid media, assays were performed by growing *S. cerevisiae* or *S. pombe* to exponential phase in YPD or YES, respectively, pelleting, and resuspending cells in water at a concentration of 0.5 OD600/mL. 4µL of 10-fold serial dilutions of cells were plated on YPD/YPEG plates or YES+glucose/YES+ethanol/glycerol and incubated at 30°C. For analysis of growth in the presence or absence of exogenous ethanolamine, cells were grown to exponential phase in SD supplemented with 10mM ethanolamine and plated on SD media or SD media supplemented with 10mM ethanolamine and incubated at 30°C.

### Whole cell extracts and Western analysis

For whole cell extracts, cells were grown to exponential phase in YPD. 0.25 OD600 cells were pelleted, washed with dH20, and extracts were prepared by alkaline extraction (0.255M NaOH, 1% 2-mercaptoethanol) followed by precipitation in 9% trichloroacetic acid. Precipitates were washed with acetone, dried, and resuspended in 50mL MURB protein sample buffer (100mM MES pH7.0, 1% SDS, 3M urea, 10% 2-mercaptoethanol) prior to Western analysis.

Protein samples were incubated at 95C for 1-2 minutes prior to SDS-PAGE, transferred to nitrocellulose membranes, and immunoblotted with *α*-Psd1 (1:1000, antibody kind gift from S. Claypool) or *α*-G6PDH (1:2000, Sigma-Aldrich A9521). Anti-rabbit antibody conjugated to DyLight 800 (1:10000, Thermo Fisher Scientific) was used and visualized with the Odyssey Infrared Imaging System (LI-COR). Linear adjustments to images were made with Photoshop 2021 (Adobe).

### Induction of LD biogenesis in S. cerevisiae, lipid isolation, and thin layer chromatography analysis

To induce LD biogenesis, cells were freshly recovered from frozen glycerol stocks and maintained in exponential growth phase in SD media for at least 16h prior to treatment with 0.5 nM *β*-estradiol. Cells were then processed for fluorescence microscopy analysis (see below) or lipid extraction and thin layer chromatography analysis. For lipid extraction, approximately 50 OD units of cells were collected for each sample, and pellet wet weight was normalized prior to extraction. Lipid extraction was performed using a modified Folch method (Folch et al., 1957). Briefly, cell pellets were resuspended in MilliQ water with glass beads and lysed by three one-minute cycles on a bead beater. Chloroform and methanol were added to the lysate to achieve a 2:1:1 chloroform:methanol:water ratio. Samples were vortexed, centrifuged to separate the organic and aqueous phases, and the organic phase was collected. Extraction was repeated a total of three times. Prior to thin layer chromatography, lipid samples were dried under a stream of argon gas and resuspended in 1:1 chloroform:methanol to a final concentration corresponding to 4 μL of solvent per 1mg cell pellet wet weight. Isolated lipids were spotted onto heated glass-backed silica gel 60 plates (Millipore Sigma 1057210001), and neutral lipids were separated in a mobile phase of 80:20:1 hexane:diethyl ether:glacial acetic acid. TLC bands were visualized by spraying dried plates with cupric acetate in 8% phosphoric acid and baking at 140°C for an hour. To quantify TLC bands, all plates were run with an internal neutral lipid standard. Densitometry of bands was performed in Fiji.

### Fluorescence microscopy and analysis

For all fluorescence microscopy analysis of *S. cerevisiae*, cells were grown in SD media with appropriate auxotrophic selection to exponential phase. To characterize LD morphology, cells were treated where indicated with 0.2% oleic acid and/or *β*-estradiol where indicated, stained with a 1:1000 dilution of MDH (AUTODOT; Abcepta) for 5 minutes, washed in water, concentrated, and immobilized directly on microscope slides. To visualize protein localization, cells were grown to exponential phase in SD media, treated where indicated with 0.2% oleic acid, concentrated, and immobilized on a 3% agarose bed of SD media on cavity microscope slides. For all fluorescence microscopy analysis of *S. pombe*, cells were grown to exponential phase in YES followed by subsequent dilution into EMM and growth for ∼16-20h before treatment/imaging as for *S. cerevisiae*.

All analysis of LD morphology was performed on a Nikon Eclipse Ti inverted epifluorescence microscope equipped with a Hamamatsu Orca-Fusion sCMOS camera and a Nikon 100x 1.45 NA objective and acquired with Nikon Elements. Z-series images were acquired with a 0.2µm step size. All images were deconvolved using AutoQuant X3 (10 iterations, blind deconvolution, low noise) and linear adjustments were made with Fiji and maximum intensity projection images are shown. All data analysis/quantification was performed on non-deconvolved (raw) images using Fiji (see below).

All analysis of subcellular protein localization was performed on a Nikon Spinning Disk Confocal microscope with Yokogawa CSU-W1 SoRa and equipped with a Hamamatsu Orca-Fusion sCMOS camera and a Nikon 100x 1.45 NA objective. Z-series images were acquired with a 0.2 µm step size and the standard resolution disk with 50 µm pinholes. Linear adjustments, except where noted in figure legends, were made with Fiji, and single plane images are shown except where noted in figures and figure legends.

For all image analysis, images were blinded prior to analysis. In experiments categorizing LD morphology, at least 75 cells were scored from at least 3 different fields of view in each of at least three independent experiments. To characterize morphology, individual cells were characterized as having normal LDs, supersized LDs (at least 0.5 µm in diameter in a single plane of view), or clustered LDs. As we determined the capability of individual cells to form supersized LDs, all cells with a mixture of morphologies were counted as supersized. In experiments quantifying LD number per cell, individual cells were scored from at least 3 fields of view imaged in each of three independent experiments.

### Electron microscopy

Yeast cells were grown to logarithmic growth phase in SD media followed by treatment with 0.2% OA for 2h and submitted to the UT Southwestern Electron Microscopy Core Facility for processing using a protocol adapted from (Wright, 2000). In brief, cells were harvested and fixed in potassium permanganate, dehydrated, and stained in uranyl acetate and embedded in Spurr Resin. Specimen blocks were polymerized at 60°C overnight and sectioned at 70 nm with a diamond knife (Diatome) on a Leica Ultracut UCT 7 ultramicrotome (Leica Microsystems). Sections were poststained with 2% uranyl acetate in water and lead citrate. Sections were placed on copper grids and poststained with 2% uranyl acetate in water and lead citrate. Images were acquired on a JEM-1400 Plus (JEOL) transmission electron microscope equipped with a LaB6 source operated at 120 kV using n AMT-BioSprint 16M CCD camera.

---

**Figure 1 – figure supplement 1.**
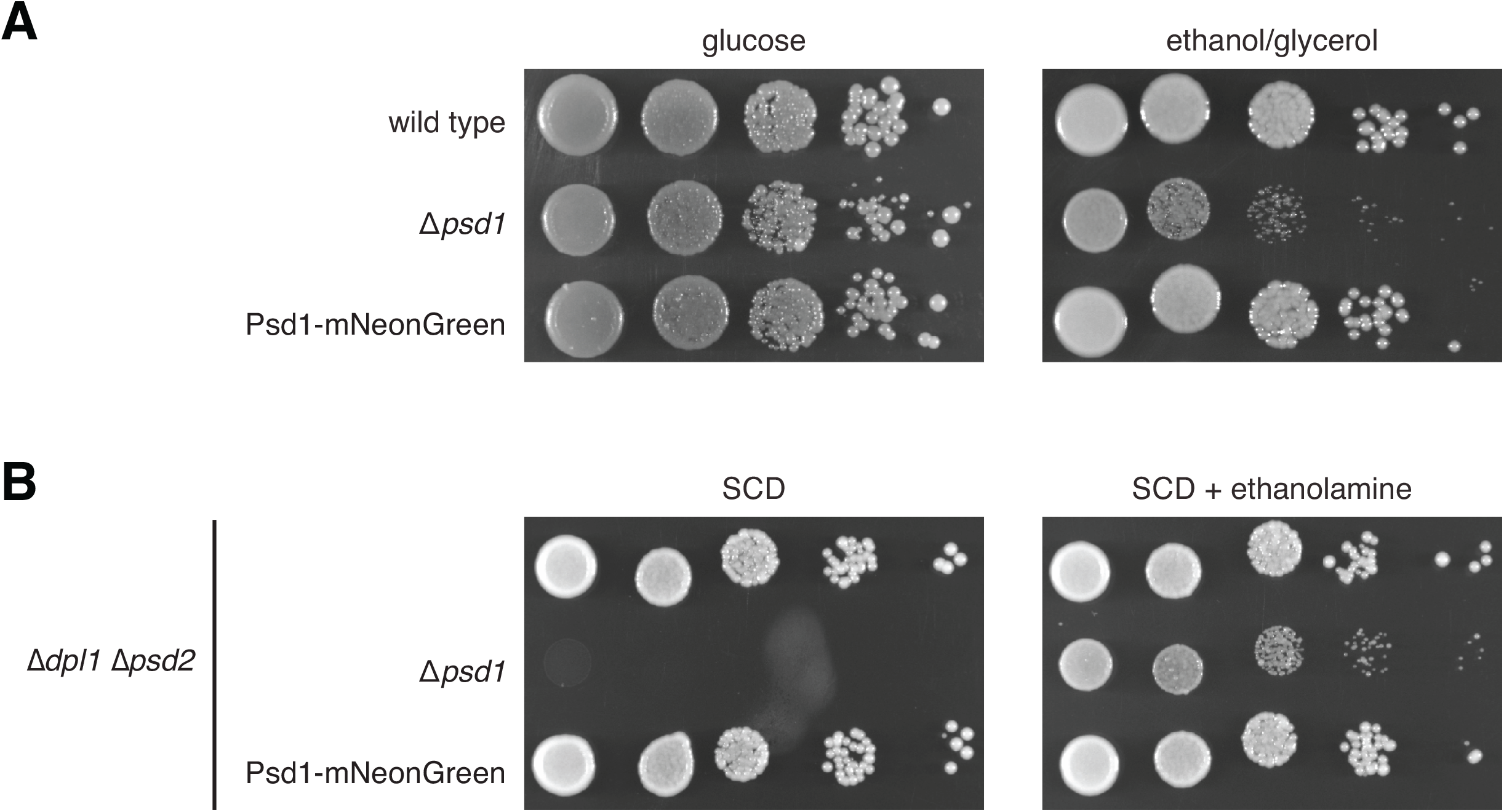
The Psd1-mNeonGreen tag is functional. **(A)** Serial dilutions of the indicated yeast cells plated on YPD media (left) or YPEG media containing the non-fermentable carbon ethanol/glycerol (right). **(B)** Serial dilutions of the indicated yeast cells plated on SD media (left) or SD media supplemented with 10mM ethanolamine (right).

**Figure 4 – figure supplement 1.**
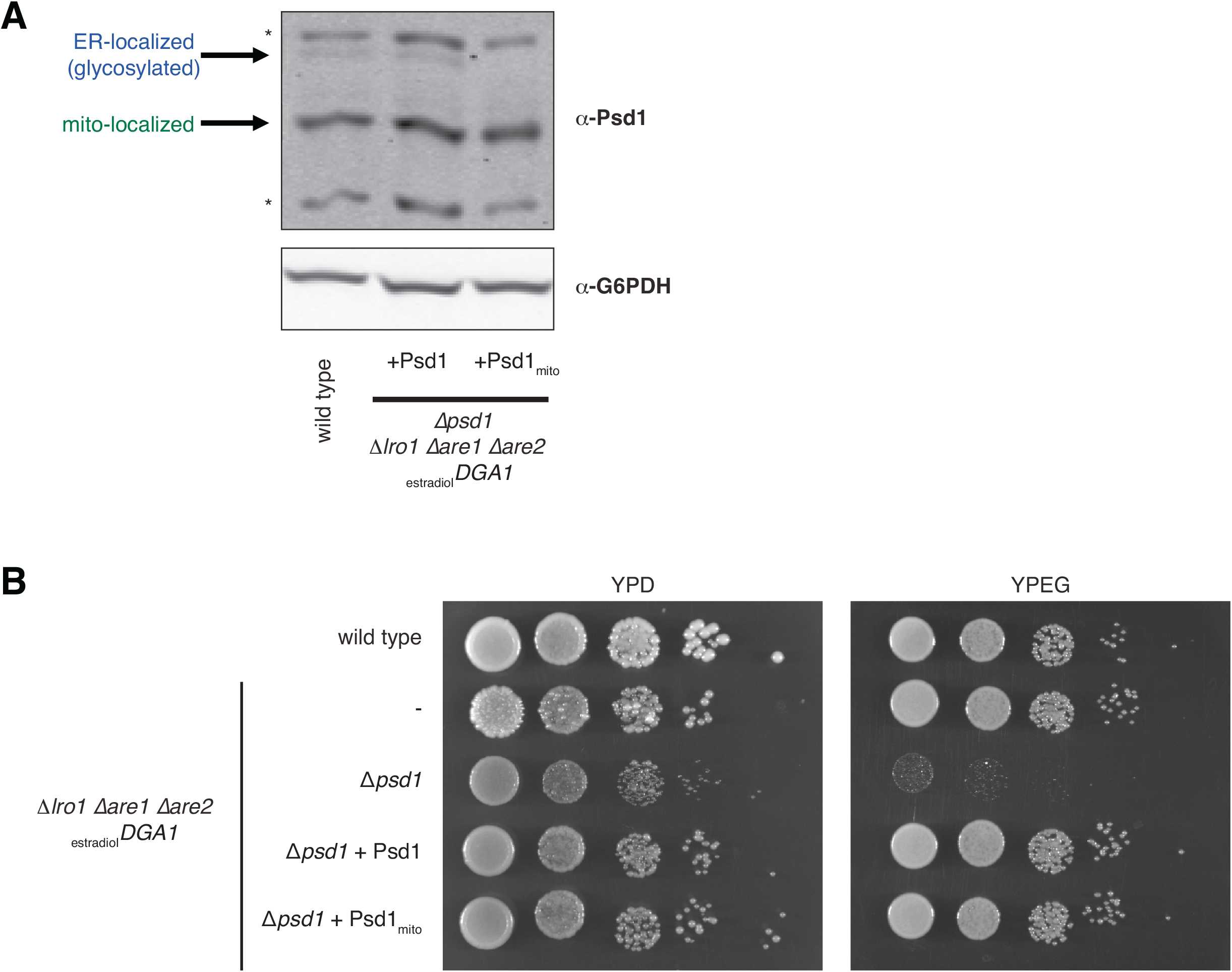
Psd1 variants reintroduced into a yeast strain with inducible LD biogenesis express and are appropriately functional. **(A)** Western analysis of whole cell lysates from the indicated strains grown in SD media with the indicated antibodies. Bands consistent with mitochondrial- and ER-localized Psd1 are indicated (Friedman et al., 2018). Asterisks mark non-specific bands. **(B)** Serial dilutions of the indicated yeast cells plated on YPD media (left) or YPEG media containing the non-fermentable carbon ethanol/glycerol (right).

